# *Phytophthora capsici* sterol reductase PcDHCR7 has a role in mycelium development and pathogenicity

**DOI:** 10.1101/2021.04.17.440084

**Authors:** Weizhen Wang, Fan Zhang, Sicong Zhang, Zhaolin Xue, Linfang Xie, Francine Govers, Xili Liu

## Abstract

The *de novo* biosynthesis of sterols is critical for eukaryotes, however, some organisms lack this pathway including most oomycetes. *Phytophthora* spp. are sterol auxotroph but remarkably, have retained a few genes encoding enzymes in the sterol biosynthesis pathway. Here we investigated the function of *PcDHCR7*, a gene in *Phytophthora capsici* predicted to encode the △7-sterol reductase. When expressed in *Saccharomyces cerevisiae*, PcDHCR7 showed a △7-sterol reductase activity. Knocking out *PcDHCR7* in *P. capsici* resulted in loss of the capacity to transform ergosterol into brassicasterol, which means PcDHCR7 has a △7-sterol reductase activity in *P. capsici* itself. This enables *P. capsici* to transform sterols recruited from the environment for better use. Biological characteristics were compared between wild-type isolate and *PcDHCR7* knock-out transformants. The results indicated that *PcDHCR7* plays a key role in mycelium development and pathogenicity of zoospores in *P. capsici*.

## Introduction

Sterols are a class of important lipids in most eukaryotes and some prokaryotes, where they may play a key role in maintaining the integrity and fluidity of cell membranes, as well as regulating biological processes (Dufourc, 2008). Sterol biosynthesis and sterol composition have been well studied in a number of organisms, and the overall picture that emerged is that the multistep biosynthesis process relies on a series of rather conserved enzymes, next to enzymes that are specific for certain lineages. As a result, the end products are different with variations in side chains and bonds of multiple rings. Within eukaryotes, fungi have ergosterol as the main sterol; vertebrates prodcuce cholesterol; *Dictyostelium* species have dictyosterol and in plants stigmasterol, campesterol, beta-sitosterol, and brassicasterol are the most common sterols (Desmond and Gribaldo, 2009). In a few prokaryotes, the sterols lanosterol and cycloartenol were identified (Pearson et al., 2003; Bode et al., 2003). For most eukaryotes, sterols are vital for their survival and it is therefore not surprising that compounds classified as sterol biosynthesis inhibitors (SBIs) have a large share in the fungicide market, in particular in agriculture for controlling fungal plant pathogens (Leaver, 2018).

Despite the vital role of sterols some organisms are sterol auxothrophs; they cannot synthesize sterols and have to recruit exogenous sterols via ingestion or absorption. Sterol auxotrophs include nematodes, most arthropods, many ciliates, and some genera of oomycetes (Shamsuzzama et al., 2020; Tomazic et al., 2014; Gaulin et al., 2010). Oomycetes are a diverse group of eukaryotic microorganisms in the Stramenopile lineage that comprises quite a number of devastating filamentous plant and animal pathogens (Govers and Gijzen, 2006; Kamoun et al., 2015). Well known oomycetes are *Phytophthora* species, important plant pathogens that cause serious damage in agriculture, forests and natural ecosystems. Examples are *Phytophthora infestans*, the causal agent of late blight and also known as the Irish potato famine pathogen (Fry, 2008), *Phytophthora ramorum*, causing sudden oak death and ravaging forests in the USA and the UK (Van Poucke et al., 2012), and *Phytophthora capsici*, a species with a wide host range and extremely devastating in many vegetable crops (Hausbeck and Lamour, 2004; Lamour et al., 2012). Although traditionally classified in the kingdom Fungi, oomycetes are phylogenetically distinct from true fungi. They evolved independently and this is also reflected at the biochemical and physiological level (Judelson and Blanco, 2005). Fungi, for example, are sensitive to SBIs; they produce large amounts of ergosterol which is essential for the plasma membrane integrity. In contrast, *Phytophthora* species are not sensitive to SBIs because they are sterol auxotrophs, and the same holds for species in the oomycete genus *Pythium*, also mainly plant pathogens, and the obligate downy mildew pathogens. For a long time, all oomycetes were thought to be sterol auxotrophs, but in the last decades this is called into question. Studies inspired by oomycete genome sequencing demonstrated that genera comprising largely animal pathogens like *Aphanomyces* and *Saprolegnia*, do possess the capacity to synthesize sterols, and, as in fungi, and some SBIs can strongly inhibit their growth (Madoui et al., 2009; Warrilow et al., 2014). It is thus conceivable that the last eukaryotic common ancestor (LECA) of oomycetes had the capacity to synthesize sterols.

Even though *Phytophthora* is sterol auxotrophic, at least two genes encoding enzymes in the sterol biosynthesis pathway were identified in different *Phytophthora* species, i.e. *ERG3* and *DHCR7* (Desmond and Gribaldo, 2009; Dahlin et al., 2017). The enzyme ERG3 is a C-5 sterol reductase while DHCR7 is a △7-sterol reductase. In *de novo* sterol biosynthesis in eukaryotes, △7-sterol reductase can remove the double bond at the seventh carbon of △7-sterols. In animals, DHCR7 is responsible for converting 7-dehydrocholesterol into cholesterol, which is the final step of cholesterol synthesis in the Kandutsch-Russell pathway (Prabhu et al., 2016). In humans, mutations in DHCR7 may lead to the Smith-Lemli-Opitz syndrome, a common recessive genetic disorder, causing amongst others, developmental defects and mental retardation (Waye et al., 2007; Prabhu et al., 2016). In plants, the homolog of DHCR7 named DWARF5, is crucial for brassinosteroid biosynthesis and, as the name implies, loss of function results in dwarfism of *Arabidopsis* (Choe et al., 2000). As yet, the function of DHCR7 in sterol auxotrophic organisms has not been studied.

The aim of this study was to investigate the function of DHCR7 in a sterol auxotrophic *Phytophthora* species. We choose *P. capsici* as model species because of its importance as plant pathogen and because it is amenable to gene editing using CRISPR/Cas9 (Miao et al., 2018). *P. capsici* causes root, crown or fruit rot in over 20 families of plants including Cucurbitaceae, Solanaceae, and Leguminosae (Silvar et al., 2006; Lamour et al., 2012). Its lifecycles can be divided into sexual and asexual ones. In the sexual lifecycle, *P. capsici* can produce long-lived dormant oospores, which can be used as the main source of preliminary infection in soil. *P. capsici* is heterothallic and thus needs two isolates with different mating types for sexual production. In contrast, in the asexual lifecycle, the branched sporangiophores emerged from hyphae can produce sporangia, and the mature sporangia can quickly release biflagellate motile zoospores that swim chemotactically and infect plants, which are the main sources for secondary infection and the spread of this disease. In this study we characterized the *P. capsici DHCR7* gene (*PcDHCR7*), analyzed its expression during the *P. capici* life cycle, and verified the enzyme activity of DHCR7 by heterologous expression in yeast. Moreover, we analyzed the phenotypes and the sterol-modifying capacity of *PcDHCR7* knock-out transformants and revealed molecular and biological functions of this gene in *P. capsici*.

## Results

### Characterization of DHCR7 and its expression profile in *P. capsici*

Although *Phytophthora* spp. are known as sterol auxotrophs, the gene *DHCR7* was found to be present in different species in this genus (Desmond and Gribaldo, 2009; Dahlin et al., 2017). Also in several *Pythium* species, which cannot synthesize sterols either, genome mining revealed the presence of a *DHCR7* homolog. Sequence alignment (Figure S1) and phylogenetic analysis (Figure 1a) showed that DHCR7 is highly conserved within oomycetes and suggest that the *DHCR7* gene was present in the last eukaryotic common ancestor (LECA) of oomycetes and before speciation. Protein signature analysis of PcDHCR7 from *P. capsici* identified six transmembrane domains (Figure 1b), indicating it is probably a transmembrane protein.

**Figure 1.**
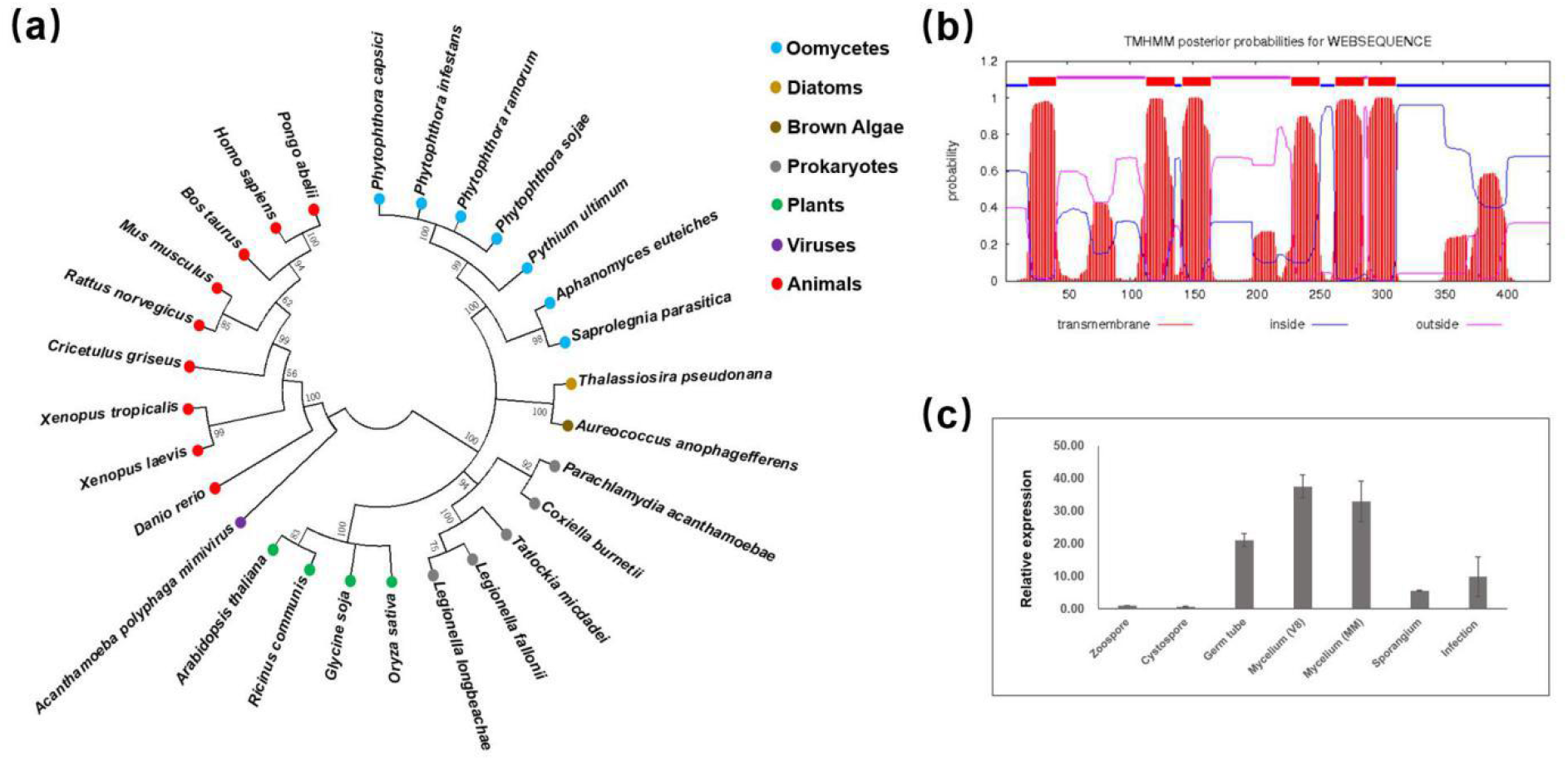
Characterization of DHCR7 protein and its expression profile in *Phytophthora capsici*. (a) Condensed molecular phylogenetic tree of DHCR7 protein sequences of representative species from different lineages. Bootstrap values are expressed as percentages based on 1000 repetitions, and only those with >50% branch support are shown. (b) Transmembrane domain analysis of PcDHCR7 using TMHMM. Six transmembrane domains are predicted to be present in the PcDHCR7 protein. (c) Expression profile of *PcDHCR7* in *P. capsici* by RT-qPCR analysis. The gene *PcDHCR7* is expressed in all development stages and during infection (4 dpi). V8 means that mycelia were cultured on V8 medium, and MM indicates that mycelia were cultured on minimal medium without any sterol. Data represent the mean ± SD from three biological repeats.

The fact that *P. capsici* cannot synthesize sterols raised the question if *PcDHCR7* is expressed and if so in which life stages and under which conditions. RNA was isolated from different life stages and from mycelium grown on minimal medium without sterols and V8 medium that is made from vegetable juice containing natural plant sterols. RT-qPCR analyses showed that *PcDHCR7* is expressed in all tested life stages, also during infection (four days after inoculation), but at different levels (Figure 1c). The *PcDHCR7* mRNA levels in germ tubes and mycelium are relatively high compared to the other life stages and there is no significant difference in expression in mycelium cultured on medium with or without sterols (Figure 1c).

### PcDHCR7 shows sterol reductase activity in *Saccharomyces cerevisiae*

To determine whether the protein encoded by *PcDHCR7* has sterol reductase activity we expressed the gene in the yeast *Saccharomyces cerevisiae*. In this model organism the sterol biosynthesis pathway is well known (Reiner et al., 2005). It lacks the DHCR7 homolog, and therefore the sterol end product—ergosterol—as well as its precursors contain a double bond at the seventh carbon. Since PcDHCR7 is a putative △7-sterol reductase it might use ergosterol and its precursors as its substrates. To test this, we cloned *PcDHCR7* in a yeast expression vector for heterologous expression in *S. cerevisiae* (Figure 2a) and analyzed the sterol composition of the transformants. The *S. cerevisiae* transformants expressing *PcDHCR7* showed a similar growth morphology and growth rate as those of control strains transformed with the empty vector (Figure 2b). Hence, the presence of PcDHCR7 protein does not impair yeast growth. After two days’ cultivation of the *S. cerevisiae* strains containing the *PcDHCR7* expressing vector as well as control strains in the liquid culture, the yeast cells were collected for sterol extraction and detection. As a result, a substantial amount of ergosterol was detected in all strains tested (Figure 2c), and brassicasterol was found in all the strains with the *PcDHCR7* expressing vector, in contrast to control strains (Figure 2c). Brassicasterol is one of the important sterols in plants, and the only difference between the structures of brassicasterol and ergosterol is the double bond at the seventh carbon, indicating that PcDHCR7 shows a △7-sterol reductase activity in *S. cerevisiae*.

**Figure 2.**
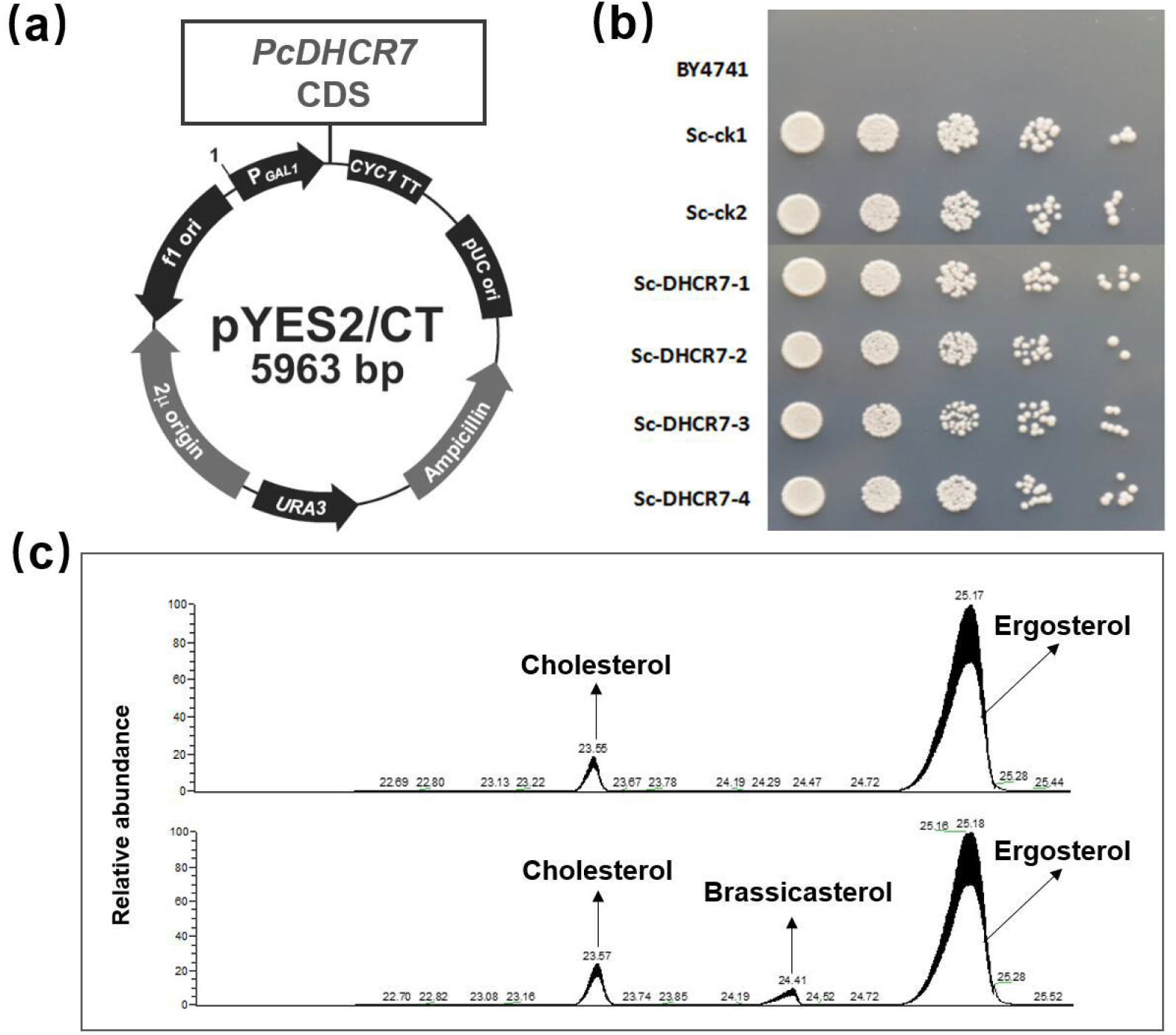
Expressing *PcDHCR7* in *Saccharomyces cerevisiae* and sterol detection from transformants. (a) The coding DNA sequence of *PcDHCR7* was inserted into the pYES2/CT yeast expression vector under the drive of the promoter GAL1, and the original vector without *PcDHCR7* was used as a negative control. (b) The growth of *S. cerevisiae* transformants and parent strain BY4741 on a selective medium (which is uracil deficient) indicates that the selective marker was expressed in transformants. Empty vector strains (Sc-ck1 and Sc-ck2) had a similar growth rate to *PcDHCR7* expressing strains (Sc-DHCR7-1, Sc-DHCR7-2, Sc-DHCR7-3, and Sc-DHCR7-4). (c) Sterols detected from the representative empty vector strain (above) and PcDHCR7-expressing strain (bottom) showed that brassicasterol was present in the latter strain. Cholesterol was used as an internal standard and was manually added during sterol extraction. All of the sterols indicated in the figure were detected as sterol derivatives with a trimethylsilyl at the C-3 hydroxyl.

### PcDHCR7 shows sterol reductase activity in *P. capsici*

For addressing the question if the endogenous *PcDHCR7* gene in *P. capsici* encodes a sterol reductase that is functional in its natural setting, we chose a loss-of-function approach taking advantage of the possibilities offered by CRISPR/Cas9 genome editing (Fang and Tyler, 2016), and obtained several homozygous knock-out transformants (Figure S2). To check the sterol reductase activity we cultured wild-type *P. capsici* and one representative *△PcDHCR7* transformant on minimal medium supplemented with ergosterol at a concentration of 20 μg/mL and subsequently extracted the sterols from the mycelium. The chromatograms for sterol detection showed major differences in sterol content between wild-type and *△PcDHCR7* transformant. Brassicasterol is the major peak in wild-type with almost no peak of ergosterol, whereas ergosterol is the only peak in *△PcDHCR7* transformant (Figure 3). These results show that *P. capsici* is capable of recruiting ergosterol from the medium and can use it as a substrate that is converted into brassicasterol. Also the *△PcDHCR7* transformant seems to recruit ergosterol from the medium but in contrast to wild-type *P. capisici*, the transformant is not capable of using ergosterol as substrate for producing brassicasterol. The only genetic difference between *△PcDHCR7* transformant and wild-type isolate is the absence of *PcDHCR7* and hence we can conclude that *PcDHCR7* is a functional gene encoding an enzyme with △7-sterol reductase activity in *P. capsici* itself.

**Figure 3.**
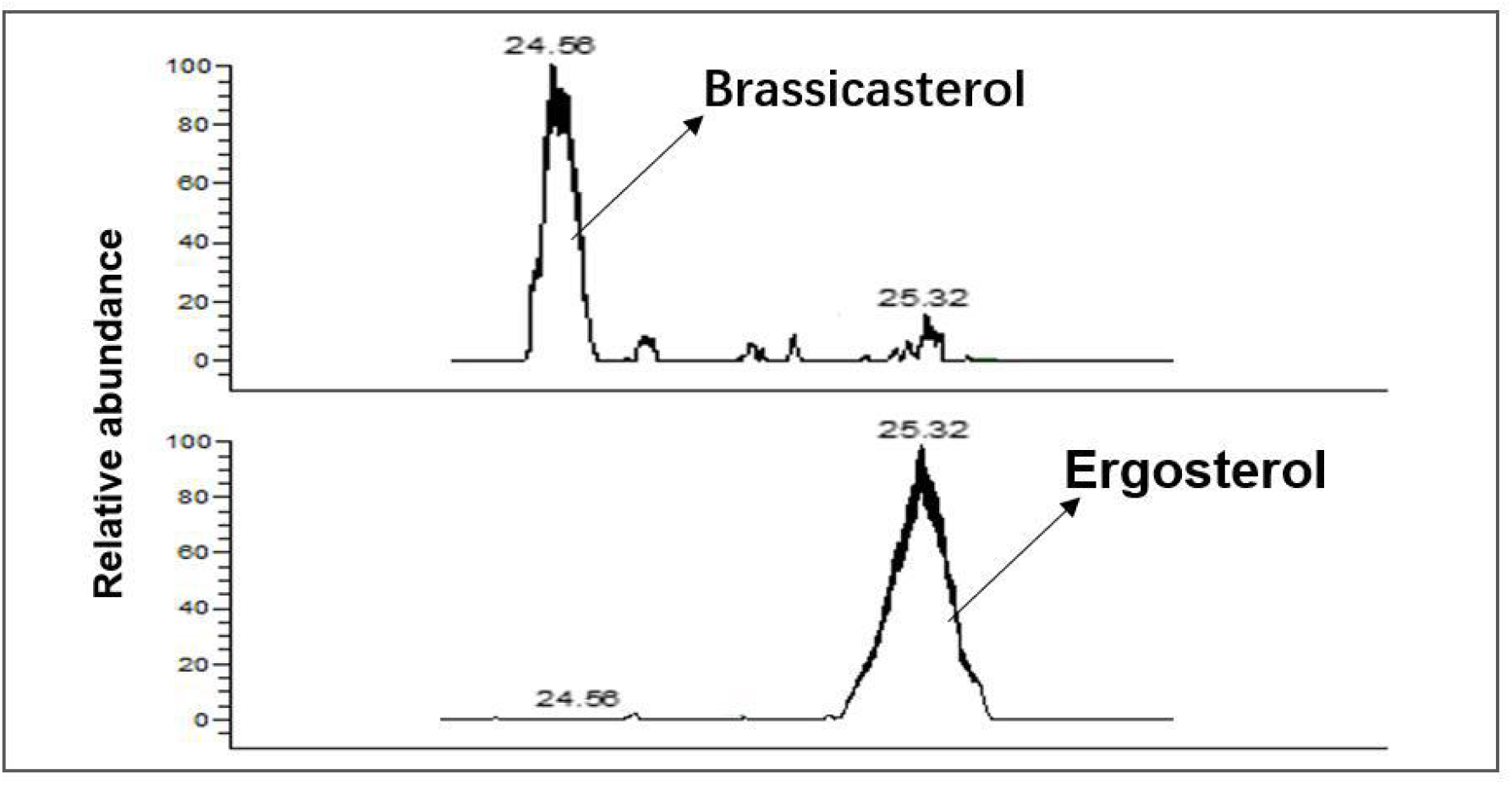
Sterol detection from different *Phytophthora capsici* isolates. Wild-type isolate BYA5 (above) and *△PcDHCR7* transformant KD1-1 (bottom) are respectively cultured on minimal medium modified with 20 μg/mL ergosterol. All of the sterols indicated in the figure were detected as sterol derivatives with a trimethylsilyl at the C-3 hydroxyl.

### Sterols saturated by PcDHCR7 promote zoospore production

Earlier studies on several *Phytophthora* spp. have shown that adding sterols to growth medium has a positive effect on growth and development (Hendrix, 1970; Marshall et al., 2001). Apparently, these sterol-auxotroph organisms can recruit exogenous sterols from the environment and utilize these for their own benefit. Here we tested the effect of sterols on asexual reproduction in *P. capsici*. On minimal medium without any sterols, *P. capsici* did not produce any zoospores, but when adding sterols zoospore production was triggered (Figure S3). All four tested sterols, i.e., ergosterol, cholesterol, β-sitosterol, and stigmasterol, promoted zoospore production in a concentration-dependent manner while reaching a plateau at 20 μg/mL (Figure S3). With the exception of ergosterol these sterols are downstream products of the biosynthesis step mediated by the △7-sterol reductase. Knowing that PcDHCR7 uses ergosterol as substrate to produce brassicasterol (as shown in Figure 3) enabled us to explore whether *P. capsici* has a preference for certain sterols. When feeding ergosterol to *P. capsici △PcDHCR7* transformant, this knock-out strain produced hardly any zoospores while the wild-type isolate showed abundant zoospore production when supplied with the same amount of ergosterol (Figure 4a). However, when replacing ergosterol by brassicasterol, both the wild-type and *△PcDHCR7* transformant produced equal amounts of zoospores (Figure 4b). This suggests that *P. capsici* has a preference for utilizing sterols that are saturated at the seventh carbon.

**Figure 4.**
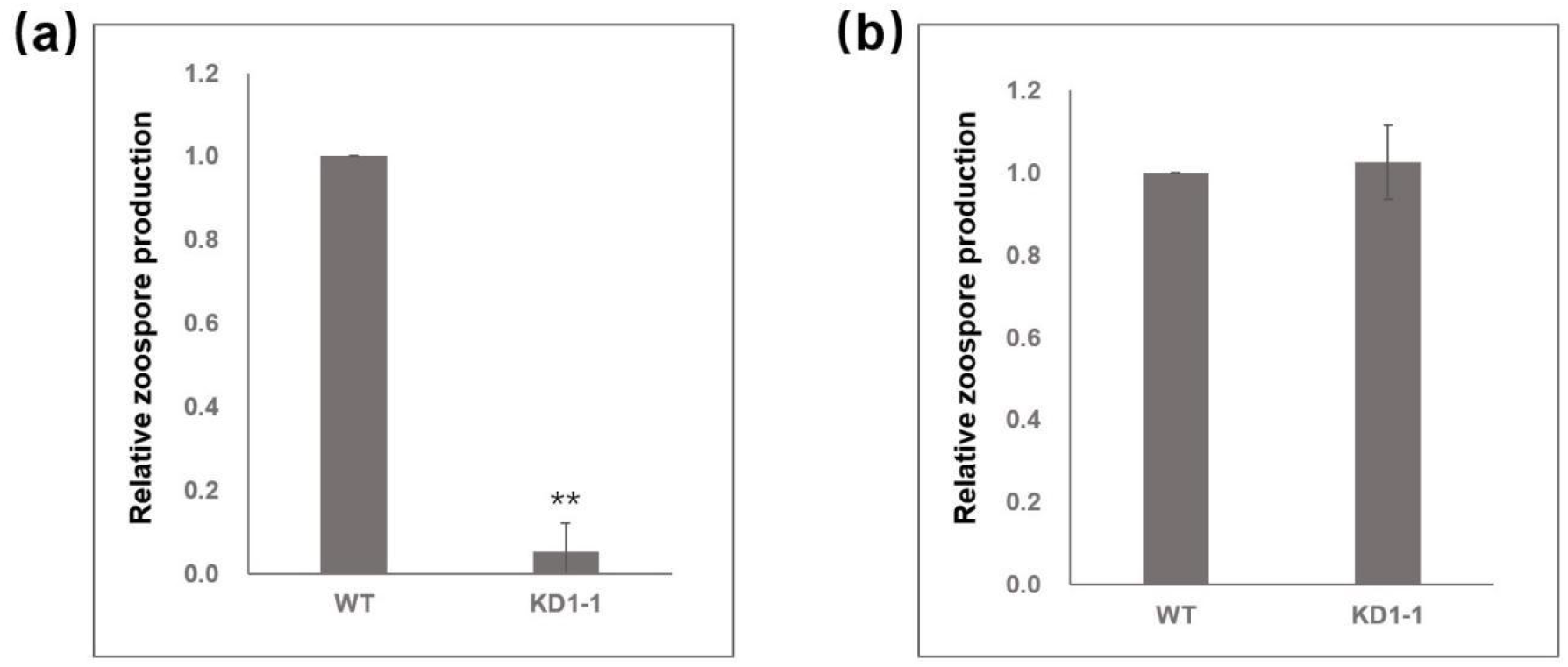
Zoospore production of *Phytophthora capsici* treated with different sterols. (a) *P. capsici* wild-type isolate and *△PcDHCR7* transformant were treated with 20 μg/mL ergosterol. (b) *P. capsici* wild-type isolate and *△PcDHCR7* transformant were treated with 20 μg/mL brassicasterol. WT indicates wild-type isolate BYA5, and KD1-1 is a representative *△PcDHCR7* transformant. Zoospore production is shown as the number of zoospores of transformant relative to that of wild-type isolate. Values represent mean ± SD of three independent experiments, and double asterisks denote significant difference from each other. **p < 0.01.

### PcDHCR7 has a role in pathogenicity of zoospores

To further investigate the role of PcDHCR7 in growth and development of *P. capsici* we analyzed mycelial growth, sporangium production, zoospore production and cystospore germination of three independent *△PcDHCR7* transformants (Figure S2). Compared to the wild-type isolate all three *△PcDHCR7* transformants showed reduced growth with significantly smaller colonies four days after inoculation on V8 agar medium (Table 1). Other phenotypes though, were not severely affected. Sporangium production, zoospore release and cyst germination rates were in the same range in the knock-out isolates and the wild-type with the exception of KD3-1 that showed a drastic reduction in sporangium production and as a consequence, also in the amount of zoospores that were released (Table 1). Based on the molecular characterization KD3-1 is a bona fide knock-out strain so why it behaves different is not clear.

**Table 1.**
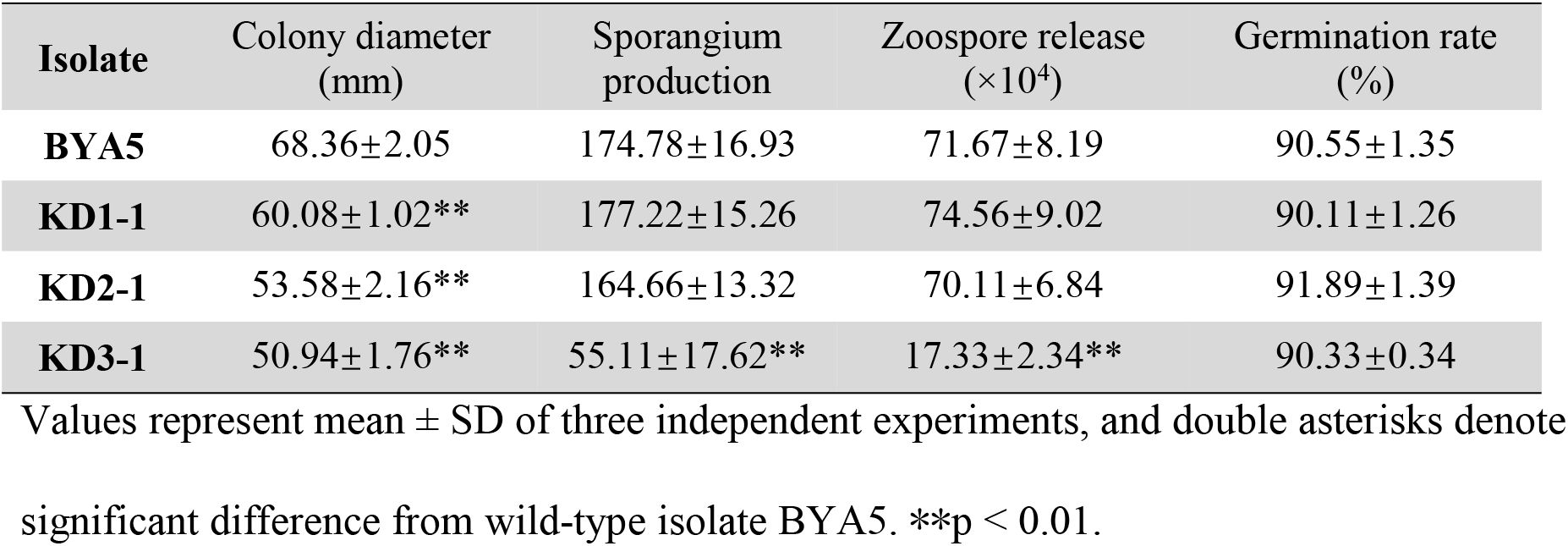
Biological characteristics of wild-type isolate and *△PcDHCR7* transformants.

Zoospore inoculation on pepper and *Nicotiana benthamiana* resulted in lesions within three days on leaves exposed to zoospores from the wild type strain; however, no lesions were visible on leaves inoculated with the knock-out strains (Figure 5a and 5d). A similar result was obtained when exposing pepper seedlings to zoospores. Five days after zoospores of the wild-type isolate were added to the potting soil the seedlings wilted but adding zoospores of the knock-out strains did not affect the health status of the plants (Figure 5b). Using a modified CRISPR/Cas9 system, the gene *PcDHCR7* was *in situ* complemented in the *△PcDHCR7* transformant KD1-1 (Wang et al., 2019). After the gene was reintroduced, the pathogenicity of zoospores was rescued partly, possibly because the expression level of the *PcDHCR7* gene in the complemented transformant was lower than that of the wild-type isolate (Figure S4). Given that the cystospores of the transformants could germinate normally compared with those of the wild-type isolate, it was inferred that the lack of zoospore infection of *△PcDHCR7* transformants was due to the invasion properties of the germ tube or developmental defects from the germ tube to the mycelium. However, cell death examination with leaves of *N. benthamiana* showed that the germ tubes of *△PcDHCR7* transformants still seemed invasive, but the infection area was only restricted in the inoculation site (Figure 5d).

**Figure 5.**
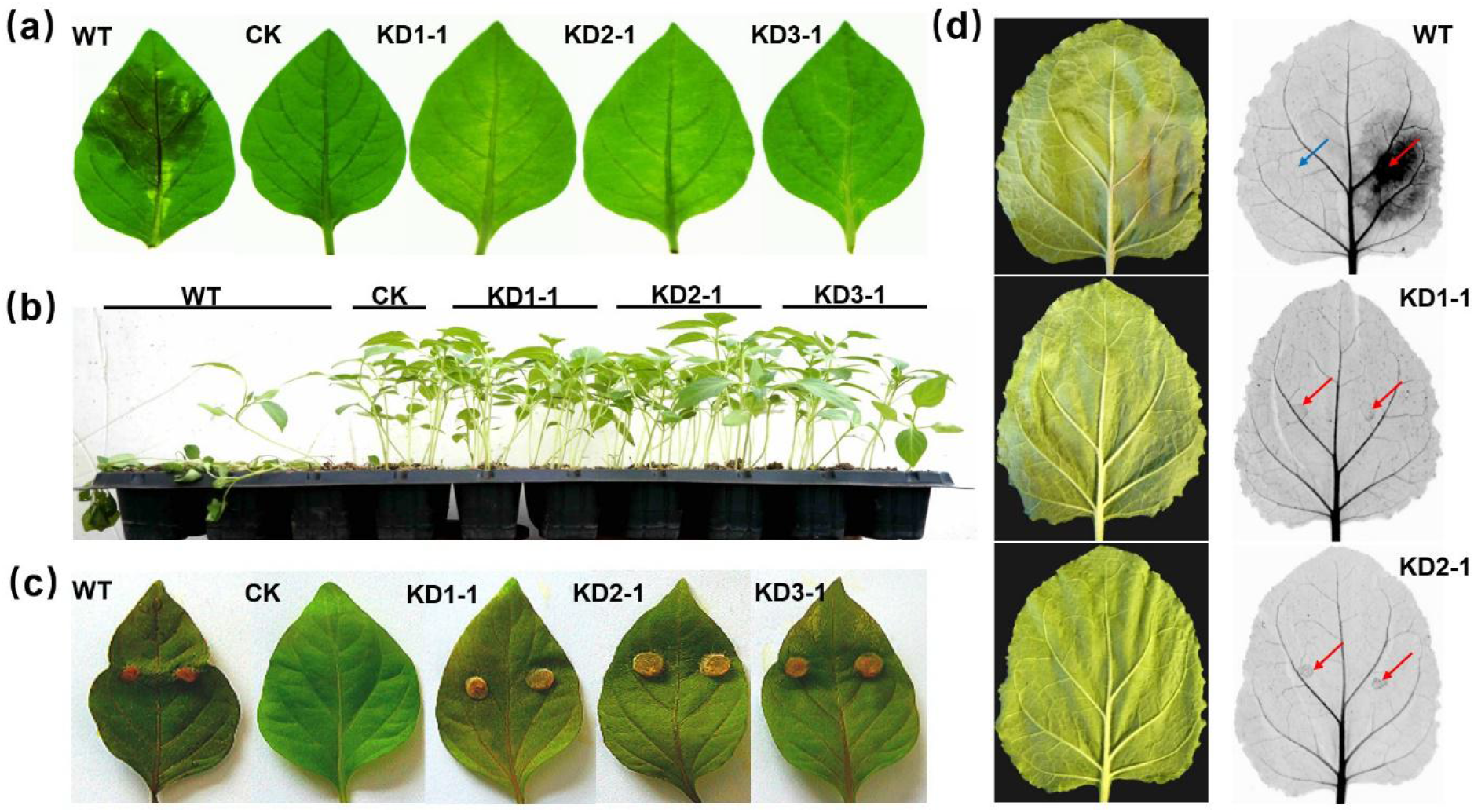
Pathogenicity evaluation of wild-type isolate and *△PcDHCR7* transformants with pepper and *Nicotiana benthamiana*. (a) Symptoms of detached pepper leaves inoculated with zoospores of different isolates (3 dpi). (b) Symptoms of seedling pepper plants inoculated with zoospores of different isolates (5 dpi). (c) Symptoms of detached pepper leaves inoculated with mycelia of different isolates (4 dpi). WT indicates wild-type isolate BYA5; KD1-1, KD2-1, and KD3-1 are representative *△PcDHCR7* transformants; CK means the leaves were treated with an equivalent amount of water, or without treatment in the case of mycelium inoculation. (d) Symptoms of detached *N. benthamiana* leaves inoculated with zoospores of different isolates (3 dpi). Blue arrow indicates inoculation site with water, and red arrow indicates inoculation site with zoospores.

### Lack of PcDHCR7 affects normal mycelium development

The finding that the encysted zoospores from the knock-out strains showed normal germination rates but nevertheless failed to cause lesions raised the question at which step in the infection process the growth of knock-out strains was hampered. To answer this, we first examined the development of the germ tube into mature mycelium under the microscope. At 24 h after zoospore encystment and germination the wild-type isolate showed elongated hyphae with occasional branching (Figure 6). In contrast, in the knock-out strains most germ tubes had stopped growing, and many hyphae were twisted with abnormal shaped tips and deformed branches (Figure 6). Complementation of the knock-out strain ReD1-1 with *PcDHCR7* almost restored the wild type morphology of the hyphae. This aberrant growth phenotype likely explains why zoospores of knock-out strains lost pathogenicity (Figure 5 a,b,d) and why the colonies of the knock-out strains had a smaller diameter when compared to the wild-type strain (Table 1). Yet, the fact that the mycelium grows, albeit slower, and that there is sporulation, suggests that the growth retardation mainly happened during the development from germ tube to mature mycelium for the △*PcDHCR7* transformants. Once that developmental phase is passed, they can grow with an almost normal pattern. To test how this affects pathogenicity we inoculated pepper leaves with mycelial plugs instead of zoospores. This head start indeed empowered the knock-out strains to cause lesions and even at a similar speed as the wild-type isolate (Figure 5c). This demonstrates that PcDHCR7 is crucial for mycelium development from germ tube during an early step in the infection process, probably after invasion at the inoculation site.

**Figure 6.**
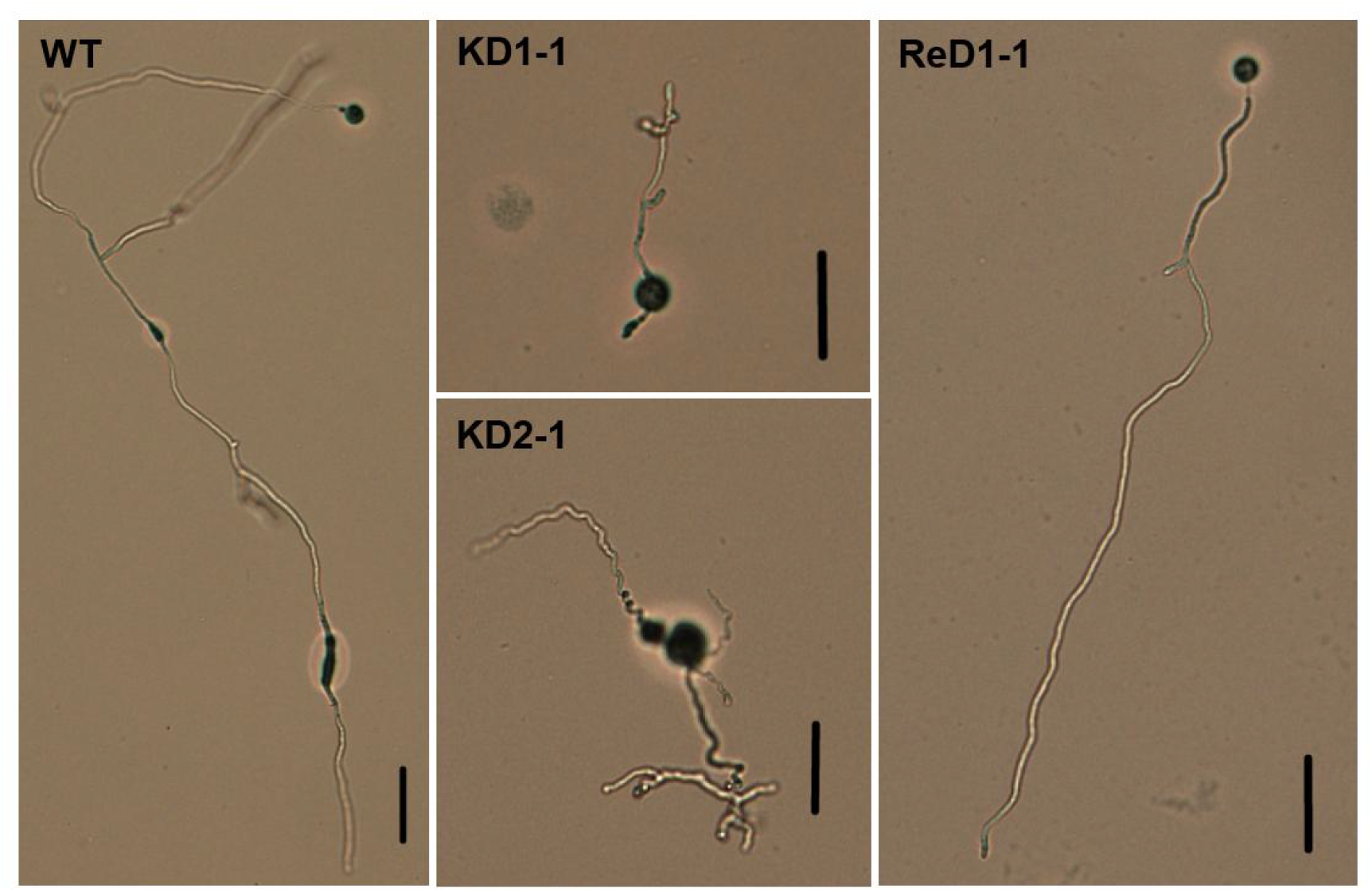
Microexamination of mycelium development of wild-type isolate and *△ PcDHCR7* transformants. Morphology of different isolates was examined after one day’s germination of cystospores. WT indicates wild-type isolate BYA5; KD1-1 and KD2-1 are representative *△PcDHCR7* transformants; ReD1-1 is a *PcDHCR7*-complemented transformant. Bar = 50 μm.

## Discussion

Sterol synthesis involves a complicated pathway in eukaryotes, and a series of sterol synthesis inhibitors targeting different proteins in this pathway of pathogens have been developed and widely used (Mercer, 1993; Müller et al., 2013). Previous studies have shown that the sterol synthesis pathway has divergently evolved in different species within oomycetes, with only a few species sustaining such an ability (Madoui et al., 2009; Warrilow et al., 2014).

Interestingly, some genes are conservatively retained in the genomes of different sterol auxotrophic oomycetes, including *Phytophthora* spp. and *Pythium* spp. This is remarkable and questions what the purpose is of maintaining these genes when the biosynthesis pathway is not functional. To address this, we investigated the enzymatic activity and the function of DHCR7 in *P. capsici*. In this study we show that *PcDHCR7* is a bona fide gene that is expressed during growth *in vitro* and *in planta* during pathogenesis, with the highest expression in mycelium. We also show that *PcDHCR7* encodes a functional enzyme that has the predicted △7-sterol reductase activity, not only in a heterologous yeast expression system, but also in *P. capsici* itself. Intriguingly, our study also revealed that PcDHCR7 is indispensable in an important phase in the life cycle of *P. capsici*. In the absence of *PcDHCR7* the transition from a germ tube to mature mycelium is severely hampered. As a consequence germ tubes emerging from encysted zoospores cannot establish a successful infection and pathogenicity is lost.

It has been shown previously that sterol auxotrophic *Phytophthora* spp. can exploit exogenous sterols for growth and reproduction (Hendrix, 1970). The effect varies depending on the sterol structures and concentrations. For example, an early study found that 10 μg/mL of sitosterol showed the best promotion of vegetative growth and sexual production for *P. sojae* (Marshall et al.,2001). The current study demonstrated that the PcDHCR7 protein has a △7-sterol reductase activity in *P. capsici* itself and can transform inactive sterols into active ones. Others showed that *P. cactorum* can transform △5,7 sterols into △5 ones (Knights and Elliott, 1976), and presumably this reduction is also mediated by DHCR7. Since *DHCR7* is conserved in all *Phytophthora* spp. analyzed so far it is likely that also the enzyme activity is maintained in other species. The question that remains is to what extent *Phytophthora* spp. are exposed to sterols that need to be transformed by DHCR7 to be profitable for the pathogen. Plants possess a broad repertoire of sterols. Moreover, concentrations and ratios vary among different plant species (Vanderplanck, 2020), and often change upon external stimulation, such as pathogen invasion (Griebel and Zeier, 2010). Even though most plant sterols have already undergone the saturation mediated by DHCR7 and lost the double bond at the seventh carbon position, it is still plausible that *Phytophthora* spp. have retained *DHCR7* in order to deal with inactive sterols recruited from their hosts under certain circumstances.

By using CRISPR/Cas9 genome editing we obtained *△PcDHCR7* knock-out strains that showed normal mycelial growth albeit with a slightly reduced growth rate. Moreover, the knock-out strains showed normal asexual sporulation and cystospore germination, indicating that *PcDHCR7* is not essential for the survival of *P. capsici*. However, the knock-out strains showed serious defects in the development from germ tube to mature mycelium and, as a result, the knock-out strains were not capable to cause lesions on plants when zoospores were used for inoculating leaves. In contrast, infection was not hampered when mycelial plugs were inoculated and this is indicative of an indirect role for DHCR7 in pathogenicity. It is likely that the aberrant growth behavior shortly after germination disables the pathogen to establish infection. The *in situ* complemented transformant used in the current study showed only half the expression level of that of the wild-type isolate, whereas a recent study observed full expression level of a knock-out gene in the *in situ* complemented *P. sojae* transformant, which encodes a regulatory B-subunit of protein phosphatase 2A (Qiu et al., 2021). This difference might result from changes in epigenetic markers on the chromatin, such as the N6-methyladenine (6mA) modification in the DNA and histone methylation, which both play a key role in regulating gene expression in *Phytophthora* (Chen et al., 2018; Wang et al., 2020). In the interplay between plants and pathogens multiple factors may play a role in sterol homeostasis in both host and pathogen. One study by Gamir et al. (2017) nicely demonstrated that a plant protein can hijack sterols in *Phytophthora brassicae*. The pathogenesis-related protein PR1, a well-known defense protein that is often upregulated upon pathogen attack, inhibits the development of *P. brassicae* in the stage from germ tube to colony. When treated with PR-1 the germ tubes displayed an abnormal morphology and the sterol hijacking seems to happen inside the cell (Gamir et al. 2017). The abnormal morphology of the *△PcDHCR7* transformants might be caused by a similar change in sterol homeostasis as caused by PR-1 via hijacking sterols. Nevertheless, the developmental defects of *△PcDHCR7* transformants might also result from malfunctioning of other pathways or other compounds that are substrates of DHCR7. Jiang et al. (2010) have used quantitative proteomics to compare embryonic brain tissue from normal mouse and a DHCR7 mutant. They found many differentially expressed proteins with putative functions in multiple biological pathways, including mevalonate metabolism, apoptosis, glycolysis, oxidative stress, protein biosynthesis, intracellular trafficking, and the cytoskeleton. Mining these data and searching for correlations and patterns might provide clues for yet unknown functions of DHCR7. It is striking that the deficiency of PcDHCR7 only influenced a very specific but short phase in the development of *P. capsici*. The mature mycelium grew happily but the stage after germination showed abnormal growth. The reason might be the difference in metabolism that produces the energy to support growth in that particular stage. In young mycelium and in the early infection stage, the main energy source is provided by fatty acid degradation, whereas in the mature mycelium, the energy is mostly produced by glycolysis, the tricarboxylic acid cycle, and the pentose phosphate pathway (Pang et al., 2017). Besides mining the omics data, more in depth analyses of the *△PcDHCR7* transformants at the cytological and biochemical level may reveal the role of this seemingly lonesome enzyme in sterol-auxotrophic *Phytophthora* spp. and this may answer the question why *Phytophthora* has retained the *DHCR7* gene.

## Methods

### Phylogenetic analysis of DHCR7 proteins

The protein sequences of DHCR7 homologs of representative evolutionary groups were acquired from different databases, including nine animals (*Homo sapiens, Pongo abelii, Bos taurus, Rattus norvegicus, Mus musculus, Cricetulus griseus, Xenopus laevis, Xenopus tropicalis*, and *Danio rerio*), four land plants (*Arabidopsis thaliana, Ricinus communis, Glycine soja*, and *Oryza sativa*), one diatom (*Thalassiosira pseudonana*), one brown algae (*Aureococcus anophagefferens*), five prokaryotes (*Legionella longbeachae, L. fallonii, Tatlockia micdadei, Parachlamydia acanthamoebae*, and *Coxiella burnetii*), one virus (*Acanthamoeba polyphaga mimivirus*), and seven oomycetes (*P. capsici, P. sojae, P. ramorum, P. infestans, Pythium ultimum, Saprolegnia parasitica*, and *Aphanomyces euteiches*). Most of the sequences were retrieved from the National Center for Biotechnology Information database (https://www.ncbi.nlm.nih.gov), except for *P. sojae, P. ramorum, P. infestans, Py. ultimum* and *Sa. parasitica* which were retrieved from the Ensembl protists database (https://protists.ensembl.org/index.html), for *P. capsici* which was retrieved from the JGI database (https://mycocosm.jgi.doe.gov/mycocosm/home), and for *A. euteiches* which was retrieved from the aphanoDB database (http://www.polebio.lrsv.ups-tlse.fr/aphanoDB/). The phylogenetic tree was constructed using Mega 6.0 (Tamura et al., 2013). The evolutionary history was inferred by using the maximum likelihood method based on the JTT matrix-based model (Jones et al., 1992). The online tool TMHMM Server v. 2.0 (http://www.cbs.dtu.dk/services/TMHMM/) was used for transmembrane domain analysis of PcDHCR7 protein.

### RNA isolation and RT-qPCR

To explore the expression profile of the *PcDHCR7* gene, biological materials from different developmental stages were collected, including zoospores, cystospores, germ tubes (about five hours after germination), 4-day-old mycelia from V8 medium, 4-day-old mycelia from minimal medium, mycelia with sporangia (four days in the dark and another five days under light), and infection stage (four days after inoculation on pepper leaves). Total RNA was extracted from the frozen samples using the SV Total RNA Isolation kit (Promega, Beijing, China), and cDNA was synthesized using the PrimeScript RT reagent Kit with gDNA Eraser (Takara, Beijing, China) according to recommended protocols. Real-time (RT)-qPCR was performed using a SYBR Premix Dimer Eraser kit (Takara, Beijing, China) on an ABI7500 sequence detection system (Applied Biosystems, United States). *Actin* and *WS21* genes were used as references for normalization of the target gene expression (Yan and Liou, 2006). The primers used for RT-qPCR are listed in Table S1. The relative expression level was calculated with the 2^-△△ CT^ method (Livak and Schmittgen, 2001), using the zoospore stage as a reference.

### PCR and multiple sequence alignments

Regular polymerase chain reactions (PCR) were conducted using a 2× master mix (Tsingke, Beijing, China), whereas those for plasmid construction were carried out using the high-fidelity DNA polymerase FastPfu system (TransGen, Beijing, China), following the recommended protocols. All of the primers used in this study are listed in Table S1. Regular PCR was performed with the following program: initial denaturing at 94 °C for 4 min, followed by 34 cycles of denaturing at 94 °C for 30 s, annealing at 55–65 °C (depending on the primer) for 30 s, and extension at 72 °C for approximately 1 min for each 1 kb fragment, with a final extension at 72 °C for 10 min. The resulting PCR products were sequenced by Tsingke (Beijing, China). Multiple sequence alignments of DNA sequences were carried out using the DNAMAN 9.0.1.116 software package (Lynnon Corporation), and alignment of amino acid sequences of DHCR7 protein from different oomycetes was performed with Clustal W (Thompson et al., 1994).

### *S. cerevisiae* transformation

The plasmid pYES2/CT was firstly digested by *EcoR*I/*Xba*I (NEB) restriction enzymes. The coding DNA sequence (CDS) of *PcDHCR7* was amplified from cDNA of the *P. capsici* wild-type isolate BYA5 and cloned into the plasmid using a modified In-Fusion® HD Cloning method (Raman and Martin, 2014). The *S. cerevisiae* strain BY4741 was used for heterologous expression of the gene *PcDHCR7*. The plasmid expressing *PcDHCR7* and the empty vector were respectively transformed into *S. cerevisiae* using the LiAc/SS carrier DNA/PEG method described in a previous study (Gietz and Schiestl, 2007) with some modifications. Yeast cells were incubated overnight in 5 mL YPD medium at 30 °C. The cultures were switched to 5–50 mL fresh YPD medium with an initial OD600 of 0.2 and incubated at 30 °C until the OD600 was approximately 1.0. The cells were collected from 5 mL cultures by centrifugation and washed with 0.1 M LiAc before they were resuspended in 26 μL sterile water, 240 μL 50% PEG (W/V), 36 μL 1.0 M LiAc, 50 mL ssDNA solution (2.0 mg/mL, going through boiling in water and cooling on ice immediately before use), and 8 μL plasmid. The mixture was placed still on ice for 5 min and heat-shocked for 40 min at 42 °C. Then, cells were collected by centrifugation, resuspended in 100 μL YPD medium, and subsequently plated on YPD agar plates without uracil. Transformation plates were placed at 30 °C for 3–4 days, and colonies were confirmed by PCR and sequencing.

### Sterol extraction and analysis

For sterol detection from *S. cerevisiae*, the yeast transformants were inoculated into liquid SD medium and incubated for 2 days for inducing expression of the target gene. Then, yeast cells were collected and washed three times with sterile water. For sterol detection from *P. capsici*, the wild-type isolate and a *△PcDHCR7* transformant were subcultured at least twice on minimal medium for *Phytophthora*; this medium does not include any sterols (Jee and Ko, 1997). Then, they were transferred to minimal medium, which was modified with 20 μg/mL ergosterol and covered with one layer of cellophane. After four days of incubation at 25 °C, the mycelia were collected. The yeast cells or *P. capsici* mycelia were dehydrated by freeze-drying, and 0.1 g dried sample was used for each detection. The cholesterol (50 μg) was added into yeast samples as an internal standard. The sterol extraction was performed using a protocol modified from a previous study (Lerksuthirat et al., 2017). After homogenization of the dried sample in 6 mL methanol-KOH (1%, w/v), saponification was conducted by heating in a water bath at 80 °C for 90 min. The sample was cooled to room temperature, then 2 mL of double-distilled water and 4 mL n-hexane were added. The mixture was homogenized overnight in a shaker and then placed still for at least 30 min before the upper organic phase was transferred into a clean tube. The solvent was evaporated in a hydroextractor, and the residue was dissolved in 200 μL methylbenzene. The N,O-bis(trimethylsilyl)-trifluoroacetamide (BSTFA, 40 μL) was added to the sample, and the derivatization reaction was carried out for 60 min at 60 °C in a water bath. For GC-MS analysis, a TSQ 8000 Evo gas chromatography in tandem with a triple quadrupole mass spectrometer was used. The sample (1 μL) was injected onto the column (Thermo TG-5MS, 0.25 μm, 0.25 mm × 30 m) with a helium flow rate of 1.2 mL/min. The temperature program was as follows: 80 °C for 1 min, followed by an increasing temperature of 12 °C/min to 280 °C (5 min) and another increasing temperature of 30 °C/min to 290 °C (5 min). The data were collected in SRM mode, and the following ions were used for the detection of different sterols: 458.5 (m/z), 353.4, and 368.4 for cholesterol; 470.5, 365.4, and 380.4 for brassicasterol; 468.5, 363.4, and 378.3 for ergosterol.

### Genome editing and growth conditions of *P. capsici*

The wild-type *P. capsici* isolate BYA5 was isolated from an infected pepper sample in Gansu Province of China in 2011. The methods used for knocking out and complementation of the *PcDHCR7* gene have been described in a previous study (Wang et al., 2019). The wild-type isolate and transformants were all maintained on solid V8 medium at 25 °C in the dark. For sterol treatment, the isolates were subcultured on minimal medium without sterols at least twice before they were transferred to minimal medium modified with different concentrations of sterols.

Sporangia and zoospores of *P. capsici* were produced based on the protocol from a previous study (Pang et al., 2013). Briefly, *P. capsici* isolates were incubated at 25 °C in the dark on solid V8 medium or minimal medium for 4 days, after which they were switched to a 12 h-light/12 h-dark photoperiod for another 6 days. The sporangium production ability was evaluated by counting all the sporangia in the entire field of vision under a microscope with 100 times magnification. The plates were flooded with 10 mL sterile water and incubated at 4 °C for 30 min and then at room temperature for 30 min. Zoospore production ability was determined by measuring zoospore concentrations using a hemocytometer. For cystospore germination evaluation, the zoospores were plated on 1% agar and dark-incubated at 25 °C for 3–8 h until most of them from the wild-type isolate had germinated. Each comparison was repeated at least three times. Single spore purification for complemented transformant was achieved by transferring individual germinated spores to fresh V8 plates, which were then dark-incubated at 25 °C for 3 days.

### Infection assays

To compare the pathogenicity of the wild-type isolate and that of transformants, zoospores and mycelia were used as inoculums, respectively. For zoospore inoculation, the concentration of different isolates was adjusted to 20,000 zoospores/mL before 10 µL zoospore suspension liquid was applied to the abaxial side of pepper or *N. benthamiana* leaves. After three or four days, the infection lesions were measured. A recently developed method using red light imaging was used for infection evaluation and cell death visualization on *N. benthamiana* leaves (Landeo Villanueva et al., 2021). On the other hand, 1 mL of zoospore suspension liquid was applied to the rhizosphere for each pepper seedling plant, which was 6–8 weeks old. After five days’ cultivation in a greenhouse, the morbidity and disease severity were investigated. For mycelium inoculation, the wild-type isolate and transformants were incubated on solid V8 medium at 25 °C in the dark, after which plugs (5 mm in diameter) with mycelia were cut from the edge of colonies and inoculated on the adaxial side of pepper leaves. After four days, the disease lesions were determined.

### Statistical Analysis

The data collected in this study were subjected to analysis of variance using DPS software ver. 9.01. Differences between means were determined using Duncan’s multiple range test at P = 0.01.

## Acknowledgments

We would like to thank Dr. Yufeng Fang and Prof. Brett Tyler (Oregon State University) for providing the original vectors for *Phytophthora* genome editing. This work was supported by the National Key Research and Development Programs of China (2017YFD0200501) and the National Natural Science Foundation of China (No. 31972304).

## Supplementary information legends

Table S1. Primers used in this study.

Figure S1. Alignment of amino acid sequences of DHCR7 protein from different oomycetes.

Figure S2. *PcDHCR7* gene knocking out in *Phytophthora capsici* and PCR confirmation of transformants.

Figure S3. Zoospore production of *Phytophthora capsici* under the treatment of sterols.

Figure S4. The pathogenicity evaluation and expression level of *PcDHCR7* in complemented transformant.

